# Expression of segment polarity genes in brachiopods supports a non-segmental ancestral role of *engrailed* for bilaterians

**DOI:** 10.1101/029892

**Authors:** Bruno C. Vellutini, Andreas Hejnol

## Abstract

The diverse and complex developmental mechanisms of segmentation have been more thoroughly studied in arthropods, vertebrates and annelids—distantly related animals considered to be segmented. Far less is known about the role of “segmentation genes” in organisms that lack a segmented body. Here we investigate the expression of the arthropod segment polarity genes *engrailed*, *wnt1* and *hedgehog* in the development of brachiopods—marine invertebrates without a subdivided trunk but closely related to the segmented annelids. We found that a stripe of *engrailed* expression demarcates the ectodermal boundary that delimits the anterior region of *Terebratalia transversa* and *Novocrania anomala* embryos. In *T. transversa*, this *engrailed* domain is abutted by a stripe of *wnt1* expression in a pattern similar to the parasegment boundaries of insects—except for the expression of *hedgehog*, which is restricted to endodermal tissues of the brachiopod embryos. We found that *pax6* and *pax2/5/8*, putative regulators of *engrailed*, also demarcate the anterior boundary in the two species, indicating these genes might be involved in the anterior patterning of brachiopod larvae. In a comparative phylogenetic context, these findings suggest that bilaterians might share an ancestral, non-segmental domain of *engrailed* expression during early embryogenesis.

## Introduction

Annelids (*e.g.* earthworms), arthropods (*e.g.* insects) and vertebrates (*e.g.* humans) are bilaterally symmetric animals that have a major part of their adult body organized into segments—sophisticated morphological units repeated along the anteroposterior axis^1^. The developmental mechanisms of segmentation are diverse and complex, but arthropods and vertebrates do share some molecular similarities in the patterning of their body segments^2–7^. These findings stimulated a debate about the evolution of segmentation that resulted in two conflicting evolutionary hypotheses. Either the similarities are the outcome of common descent, and thus support the homology of bilaterian body segments^2,8–10^, or they represent evolutionary convergences^11–13^.

These hypotheses, however, are based on the examination of distantly related groups. Annelids, arthropods and vertebrates belong to distinct branches of the Bilateria, Spiralia, Ecdysozoa and Deuterostomia, respectively (Figure 1a). This sole comparison over long evolutionary distance can be misleading to reconstruct the evolution of segmentation, because the ancestral conditions within more closely related taxa remain unknown. Thus, distinguishing homology from convergence requires a robust phylogenetic framework and dense taxonomic sampling^11,14^. To better understand the evolution of segmentation, it is necessary to investigate the related groups of annelids, arthropods and vertebrates. Most of these lineages do not have body segments, but can display a variety of serially repeated structures, such as neural ganglia, excretory organs, coeloms, and others^1,12,15^. Since segmentation is not an “all-or-nothing” trait^15,16^, further data on the morphology and gene networks in these lineages can help to elucidate the evolution of the developmental mechanisms of segmentation.

**Figure 1.**
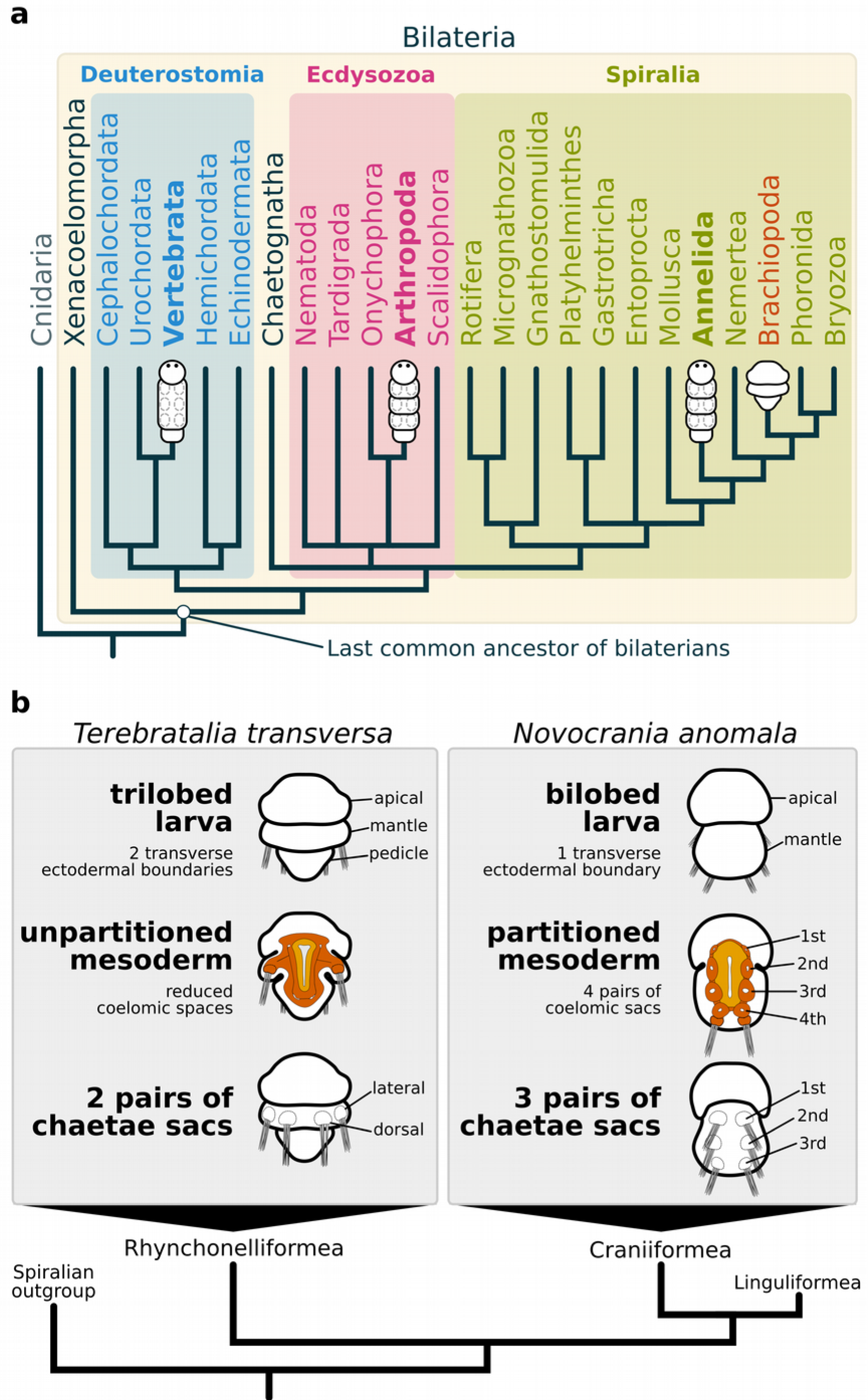
Phylogenetic relationships between bilaterally symmetric animals and the morphological features of larval brachiopods. (a) Phylogenetic tree based on recent data^29,60,61,74^ highlighting the three major bilaterian clades, Deuterostomia, Ecdysozoa and Spiralia. Groups traditionally regarded as truly segmented are marked in bold. Brachiopoda is highlighted in orange. (b) Ectodermal and mesodermal larval traits of the brachiopods *T. transversa* and *N. anomala*, as representatives of the Rhynchonelliformea and Craniiformea^47^.

A crucial mechanism of segmentation is the ability to establish tissue boundaries during animal development^17^. One of the best studied developmental boundaries are the parasegmental boundaries of the fruit fly *Drosophila melanogaster*. Fly segmentation occurs by the simultaneous division of the trunk into fourteen parasegments, developmental regions that lie out of register with the larval segments^18^. At the molecular level, the parasegment boundary is established by the abutting domains of *engrailed* (*en*) and *wingless* (*wnt1*)^19^, and is maintained by a positive feedback loop mediated by *hedgehog* (*hh*)^20^. Therefore, *en*, *wnt1* and *hh* are commonly referred to as segment polarity genes^21–23^, and their expression and function is conserved among the body segments of other arthropods^7,24–26^. The expression of segment polarity genes in annelids is more variable and, in general, do not suggest a role in the patterning of body segments^5,27^. So far, the only exception is the annelid *Platynereis dumerilii*, where the expression of *en*, *wnt1* and *hh* correlates with the segmental boundaries^10,28^. Thus, to better understand the ancestral developmental functions of the segment polarity genes in the spiralians, it is important to investigate gene expression in groups more closely related to annelids.

Brachiopods are bivalved marine invertebrates with close relations to other lophophorates, nemerteans, annelids and molluscs^29^ (Figure 1a). Even though adult brachiopods do not have body segments, the larval stages show putative repeated structures in ectodermal and mesodermal tissues. Externally, brachiopod larvae exhibit two to four lobes along the anteroposterior axis that are divided by transverse ectodermal boundaries. These lobes were once homologized to annelid segments^30^, but the observation that the ectodermal boundaries do not involve the underlying mesoderm weakened this hypothesis^31^. The mesoderm morphology is variable between brachiopod species, it can be partitioned into two or four pairs of coelomic sacs^32,33^ or not subdivided. The discovery of a brachiopod larvae with serially arranged coelomic sacs^33^ has revived the idea that brachiopods had a segmented ancestor^34,35^. Therefore, these putative segmented structures, and the closer phylogenetic position to annelids, place brachiopods as an interesting group to test the involvement of the segment polarity genes in other developmental boundaries, and to better comprehend the evolution of segmentation mechanisms in protostomes.

To investigate the developmental and molecular features of the boundaries in brachiopod embryos, we studied the trilobed larva of *Terebratalia transversa*^36^, and the bilobed larva of *Novocrania anomala*^33,37^, species that belong to distinct brachiopod lineages (Figure 1b). In these two species, we analyzed the expression of the segment polarity genes *en*, *wnt1*, and the core components of the Hedgehog signaling pathway to test whether their expression correlate with the development of the ectodermal and mesodermal boundaries of *T. transversa* and *N. anomala* larvae. Furthermore, we examined upstream control genes of *en* and discovered similarities between the molecular profile of a brachiopod larval boundary and the embryonic patterning of anterior boundaries in deuterostomes, such as the hemichordate collar/trunk boundary and the vertebrate fore/midbrain boundary, further suggesting a non-segmental ancestral role of *en* for bilaterians.

## Results

### Ectodermal and mesodermal boundaries of larval brachiopods

In *T. transversa*, gastrulation occurs by invagination and results in a radially symmetric gastrula that elongates in the anteroposterior axis (Figure 2a–c). A transverse ectodermal furrow (indentation of the epidermis) appears above the midline of the bilateral gastrula stage, extending from the blastopore lips to the dorsal side (Figure 2c), with epidermal cells posterior to the indentation being more elongated (Supplementary Fig. S1). The ectodermal furrow and cell shape differences at the bilateral gastrula are the first morphological manifestation of the boundary between the future apical and mantle lobes in *T. transversa* (apical/mantle boundary). At subsequent developmental stages, this furrow deepens and clearly divides the apical lobe from the remainder of the embryo (Figure 2d).

**Figure 2.**
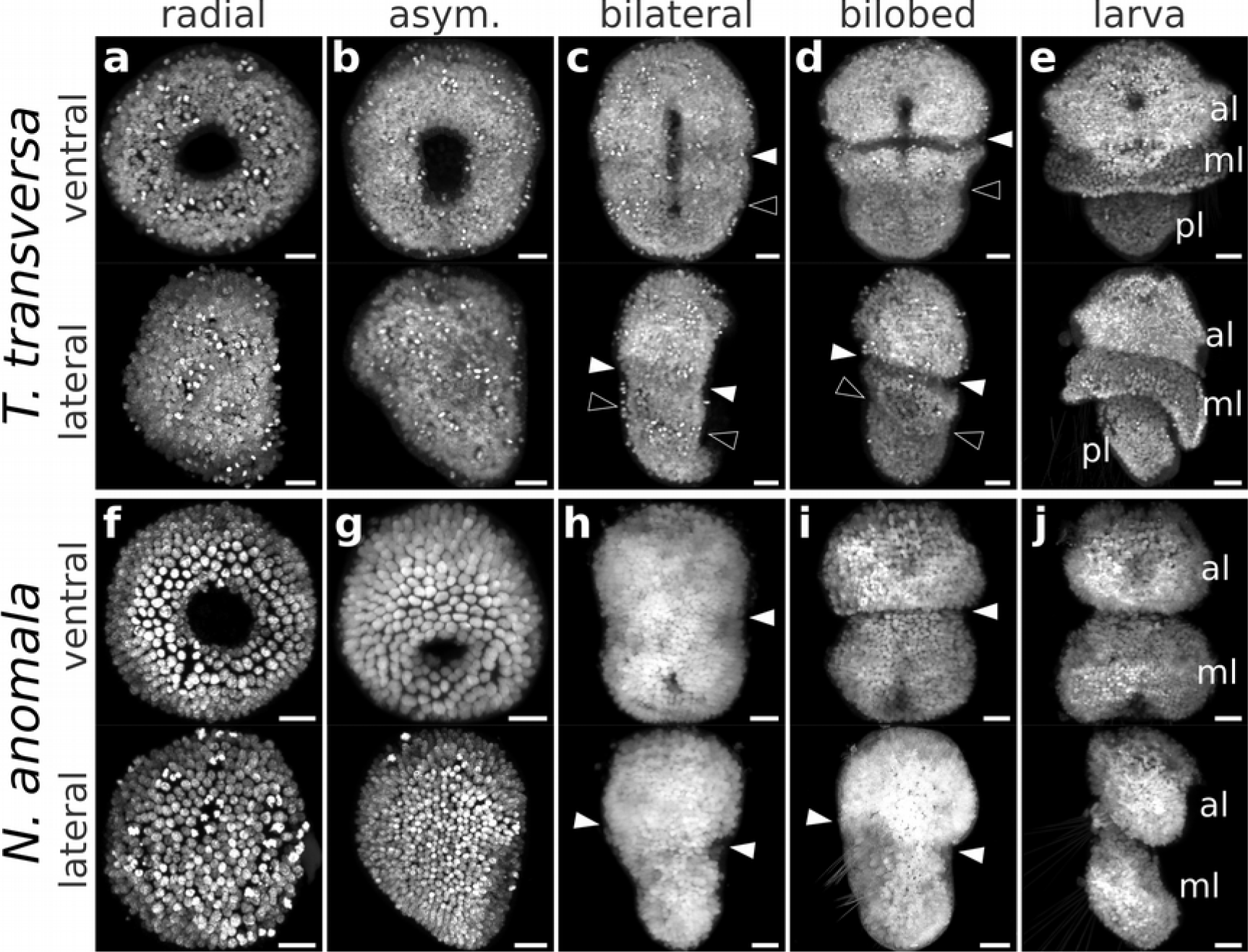
Developmental stages of the trilobed larva of *T. transversa* (a–e) and the bilobed larva of *N. anomala* (f–j). Each panel shows a ventral view (top) and lateral view (bottom) of a maximum intensity projection of embryos stained with DAPI. Anterior is top in all panels and ventral is to the right in all lateral views. White arrowheads mark the apical/mantle boundary and black arrowheads mark the mantle/pedicle boundary. al: apical lobe, ml: mantle lobe, pl: pedicle lobe. Scale bars = 20 µm.

*T. transversa* exhibits a third lobe at the posterior end, the pedicle lobe (Figure 2e). The boundary between the mantle and pedicle lobes (mantle/pedicle boundary) is identified by the narrowing of the posterior portion of the embryo (Figure 2d) and by the subsequent outgrowth of the mantle lobe (Figure 2e). The morphology of the mantle/pedicle boundary differs from the apical/mantle boundary because it is demarcated by an ectodermal fold, rather than by an indentation furrow. For this reason, the ectodermal boundaries of the trilobed brachiopod larva are not repeated, but unique structures of the larval body. In *N. anomala*, the apical and mantle lobes are also demarcated by an ectodermal furrow, as seen in *T. transversa* (Figure 2f–i), but its larva does not form a pedicle lobe (Figure 2i). Thus, the morphology of the apical/mantle boundary is conserved between both species of brachiopods.

The mesoderm of *T. transversa* has prominent projections associated to the chaetae sacs in the mantle lobe, but there are no partitions (Supplementary Fig. S2). In contrast, the mesoderm of *N. anomala* has lateral constrictions that individualize four pairs of coelomic sacs, but the posterior pouches are also tightly associated to the three pairs of dorsal chaetae sacs, while remaining interconnected more medially in the ventral side (Supplementary Fig. S2).

### Expression of the segment polarity genes *hh*, *en* and *wnt1*

To compare the ectodermal and mesodermal boundaries of brachiopods with the segment boundaries of arthropods and annelids, we analyze the expression of the segment polarity genes *en*, *wnt1* and *hh* during the embryonic development of *T. transversa* and *N. anomala*. Expression of *hh* localizes to the blastoporal lip and invaginating endomesoderm during early gastrulation, and is restricted to the endoderm in later stages (Figures 3a and 4a). In *N. anomala*, we detected an additional transverse ventral domain of *hh* near the animal pole that disappears in the bilobed stage (Figure 4a). Since the Hedgehog ligand can signal across embryonic layers, we further analyzed the expression of the Hedgehog receptors *ptc* and *smo* and the transcription factor *gli*. Transcripts are expressed in the mesoderm of *T. transversa* and *N. anomala*, as well as in the ectodermal apical and mantle territories (Supplementary Figs. S3 and S4). Altogether, the expression of *hh* and downstream pathway components does not correlate spatially or temporally with the development of the larval ectodermal boundaries of *T. transversa* and *N. anomala*.

**Figure 3.**
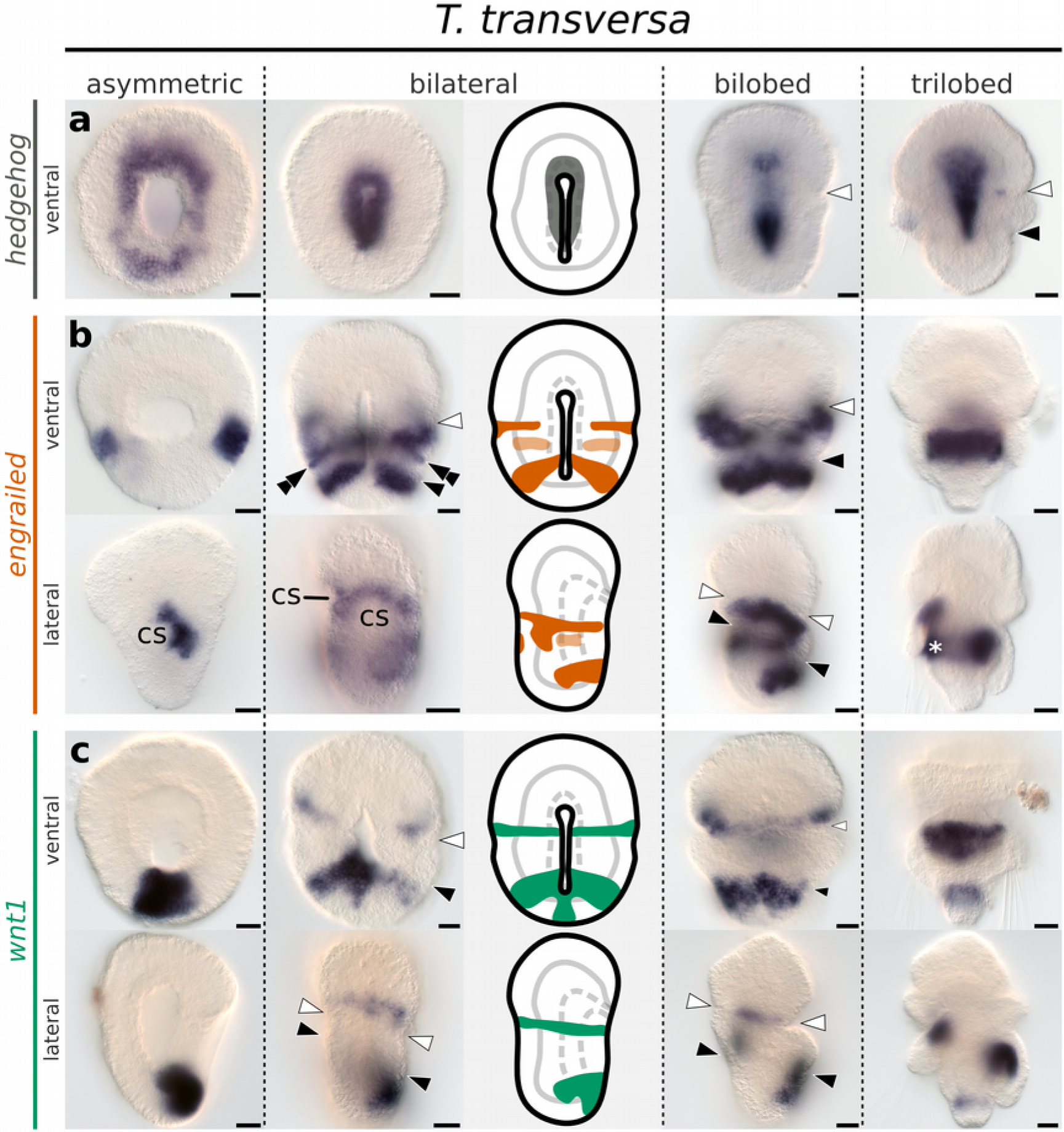
Whole mount in situ hybridization of *hh*, *en* and *wnt1* orthologs in representative developmental stages of the brachiopod *T. transversa*. Anterior is top in all panels and ventral is to the right in all lateral views. (a) Expression of *hh*. (b) Expression of *en*. (c) Expression of *wnt1*. Line drawings represent expression at the bilateral gastrula stage. White arrowheads mark the apical/mantle boundary and black arrowheads mark the mantle/pedicle boundary. Double arrowheads mark the ectodermal expression of *en*. Asterisk marks area of unspecific staining due to the rudiment of the larval shell (b, trilobed). cs: chaetae sacs. Scale bars = 20 µm.

**Figure 4.**
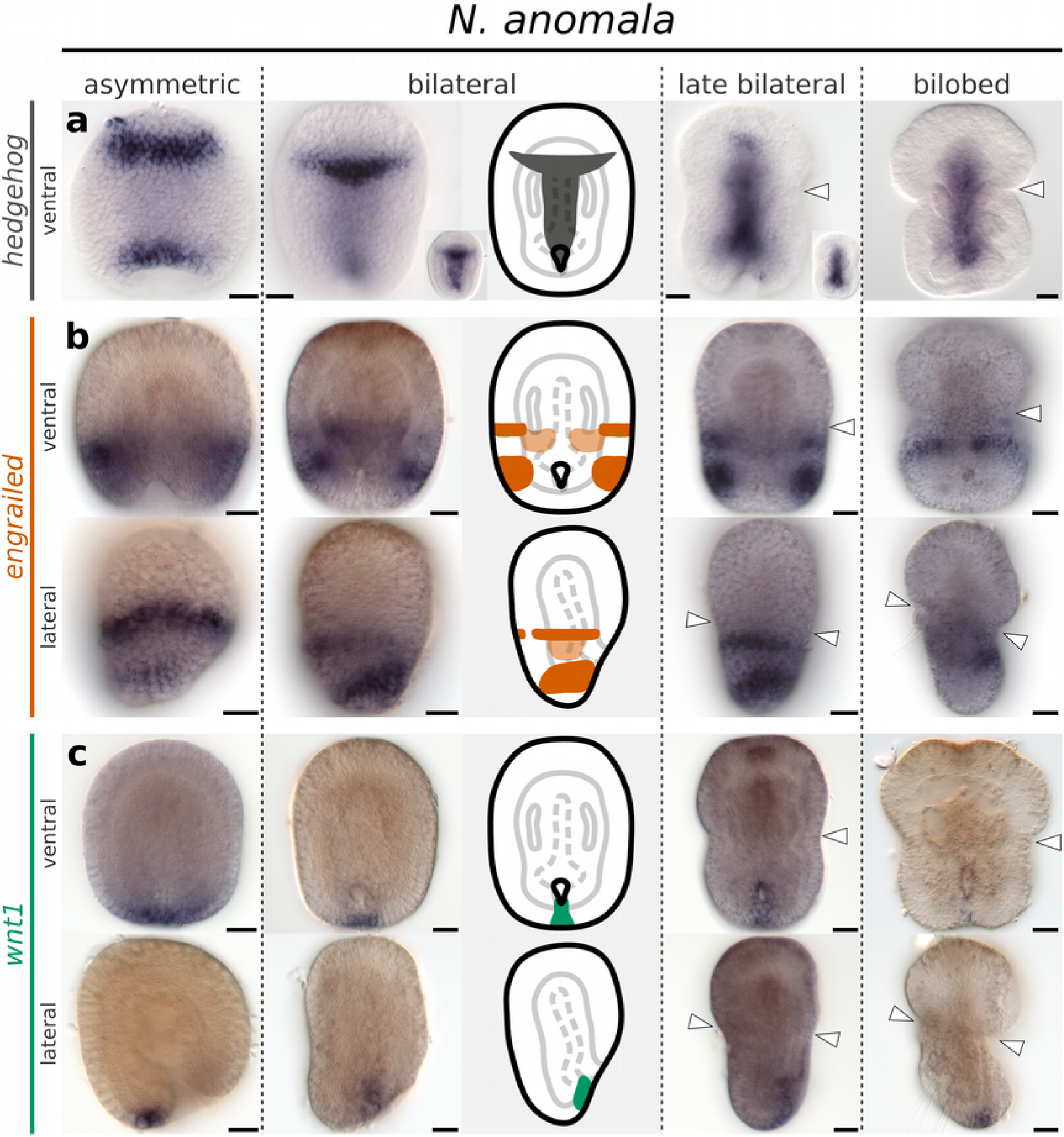
Whole mount in situ hybridization of *hh*, *en* and *wnt1* orthologs in representative developmental stages of the brachiopod *N. anomala*. Anterior is top in all panels and ventral is to the right in all lateral views. (a) Expression of *hh*. (b) Expression of *en*. (c) Expression of *wnt1*. Line drawings represent expression at the bilateral gastrula stage. White arrowheads mark the apical/mantle boundary. Scale bars = 20 µm.

We first detect transcripts of *en* in *T. transversa* in the radial gastrula stage. The expression forms an almost-complete ring in the midregion of the embryo and, progressively fades from the anterior and posterior end (Supplementary Fig. S5), leaving a pair of lateral domains of *en* in the asymmetric gastrula stage (Figure 3b). These domains extend ventrally and dorsally without reaching the blastopore on the ventral side of the bilateral gastrula stage (Figure 3b). Transcripts of *en* border the apical/mantle furrow in the subsequent stage and fade in the trilobed larvae (Figure 3b). In *N. anomala* a single dorsal ectodermal stripe of *en* expression occurs in the radial gastrula (Supplementary Fig. S5) and a second stripe emerges at the posterior end of the asymmetric gastrula (Figure 4e). Similar to *T. transversa*, *en* stripes extend ventrally without reaching the embryo midline and fade in the developed larva (Figure 4e). Therefore, expression of *en* is consistent between the two species, with the apical/mantle furrow forming immediately anterior to a pair of *en* stripes.

In addition to the striped domains demarcating the apical/mantle boundary, we identified two more posterior *en* territories on the ventral and dorsal regions of larval brachiopods, localized in the anterior-most region of the pedicle lobe. One is located on the ventral side of *T. transversa* and consists of a pair of *en* ectodermal bands associated with the blastopore in the bilateral gastrula stage (Figure 3b). The other is located on the opposite side, at the center of the dorsal surface, as a triangular-shaped domain of *en* expression (Supplementary Fig. S5). A correspondent domain also occurs on the dorsal side of *N. anomala* and localizes to the shell rudiment in the bilobed larva stage (Supplementary Fig. S5).

Mesodermal expression of *en* initiates later than the ectodermal domains. In *T. transversa*, transcripts of *en* form a pair of bands in the pedicle mesoderm that are connected to the ectodermal domains, while the mesodermal expression in *N. anomala* localizes directly inner to the lateral ectodermal domains of *en* (Supplementary Fig. S6). Expression of *en* is located to the posterior portion of the second and third pair of coelomic sacs, but not in the first or fourth (Supplementary Fig. S6). Overall, the mesodermal expression of *en* occurs in contiguity with the preceding ectodermal domains in both species.

Expression of *wnt1* in the asymmetric gastrula of *T. transversa* is associated with the posterior portion of the blastopore (Figure 3c) and shows no spatial correlation to the lateral *en* domains (Figure 5a). At the bilateral gastrula, a transverse pair of stripes of *wnt1* expression appears in the apical lobe of *T. transversa*, bordering the apical/mantle furrow anteriorly (Figure 3c). The expression abuts *en* at the apical/mantle boundary without overlap (Figure 5b). As the apical/mantle furrow deepens in the bilobed larva, *wnt1* and *en* transcripts resolve into well-defined non-overlapping stripes in *T. transversa* (Figure 5c). The stripes demarcate precisely the morphological furrow at the apical/mantle boundary of *T. transversa* with *wnt1* positioned anterior and *en* positioned posterior to the furrow (Figure 5c). In contrast, the posterior domain of *wnt1* develops a tight expression overlap with the pedicle lobe domain of *en*, on the ventral side of *T. transversa* (Figure 5b–c). Such precise coexpression pattern also occurs on the dorsal side, when *wnt1* transcripts are first detected within the triangle-shaped territory of *en* (Figure 5c).

**Figure 5.**
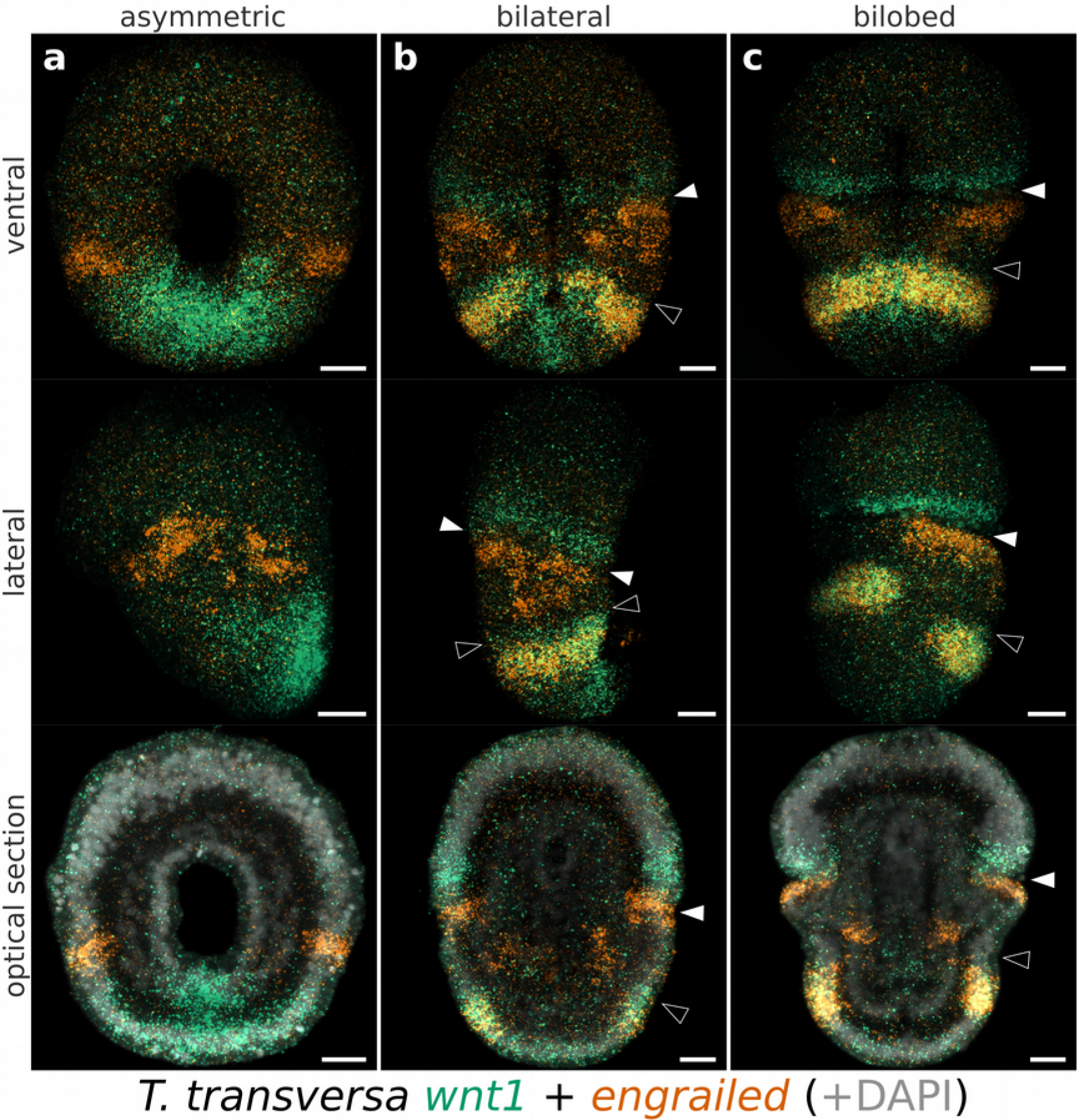
Whole mount double fluorescent in situ hybridization with *en* (vermillion) and *wnt1* (green) in *T. transversa*. (a) Asymmetric gastrula. Same specimens of Figure 2b. (b) Bilateral gastrula. Same specimens of Figure 2c. (c) Bilobed larva. Ventral view is the same specimen of Figure 2d. Ventral and lateral views are maximum intensity projections. Optical sections show additional counter staining with DAPI (gray). Merge (yellow) highlights the areas of *en* and *wnt1* co-expression. Scale bars = 20 µm.

*N. anomala* does not exhibit *wnt1* domains in the apical lobe. The transcripts are exclusively at the posterior end, associated with the blastopore (Figure 4c). We did not detect any coexpression sites for *en* and *wnt1* in *N. anomala*. Thus, the expression of *wnt1* at the apical/mantle boundary is not consistent between brachiopods, and only *T. transversa* has *wnt1* stripes demarcating the morphology of the furrow.

### Expression of the putative *engrailed* regulators *pax6*, *pax2/5/8* and *fgf8/17/18*

The only gene consistently related to the apical/mantle boundary in both brachiopod larvae is *en*. Its early expression, however, does not encircle the circumference of the embryo (Figure 3b), suggesting that there must be upstream factors positioning the apical/mantle boundary. Regulators of *en* known from *D. melanogaster* segmentation, such as *even-skipped* or *sloppy paired*^38,39^, do not correlate to the *en* domains at the apical/mantle boundary of *T. transversa*^40^. Therefore, we selected further candidate genes related to the patterning of the vertebrate neuroectoderm, where *en* plays a crucial role in the establishment of boundaries^41^. In the mid/hindbrain boundary, *en* is upregulated by *pax2* in a positive feedback loop, represses and is repressed by *pax6*, and, together with *fgf8*, is essential to maintain the boundary between the diencephalon and mesencephalon^42–45^. Thus, we analyzed the expression of the correspondent brachiopod orthologs *pax6*, *pax2/5/8* and *fgf8/17/18* in relation to the expression of *en* and the development of the apical/mantle boundary.

Transcripts of *pax6* form a pair of broad anterior domains that occupy the entire anterior half of *T. transversa* radial gastrula, and localize to the apical lobe in subsequent stages (Figure 6a). In *N. anomala*, *pax6* expression initiates in the radial gastrula as an apical ring and also localizes to the apical lobe in later stages (Figure 7a). In both species the posterior limit of *pax6* expression borders the apical/mantle furrow.

**Figure 6.**
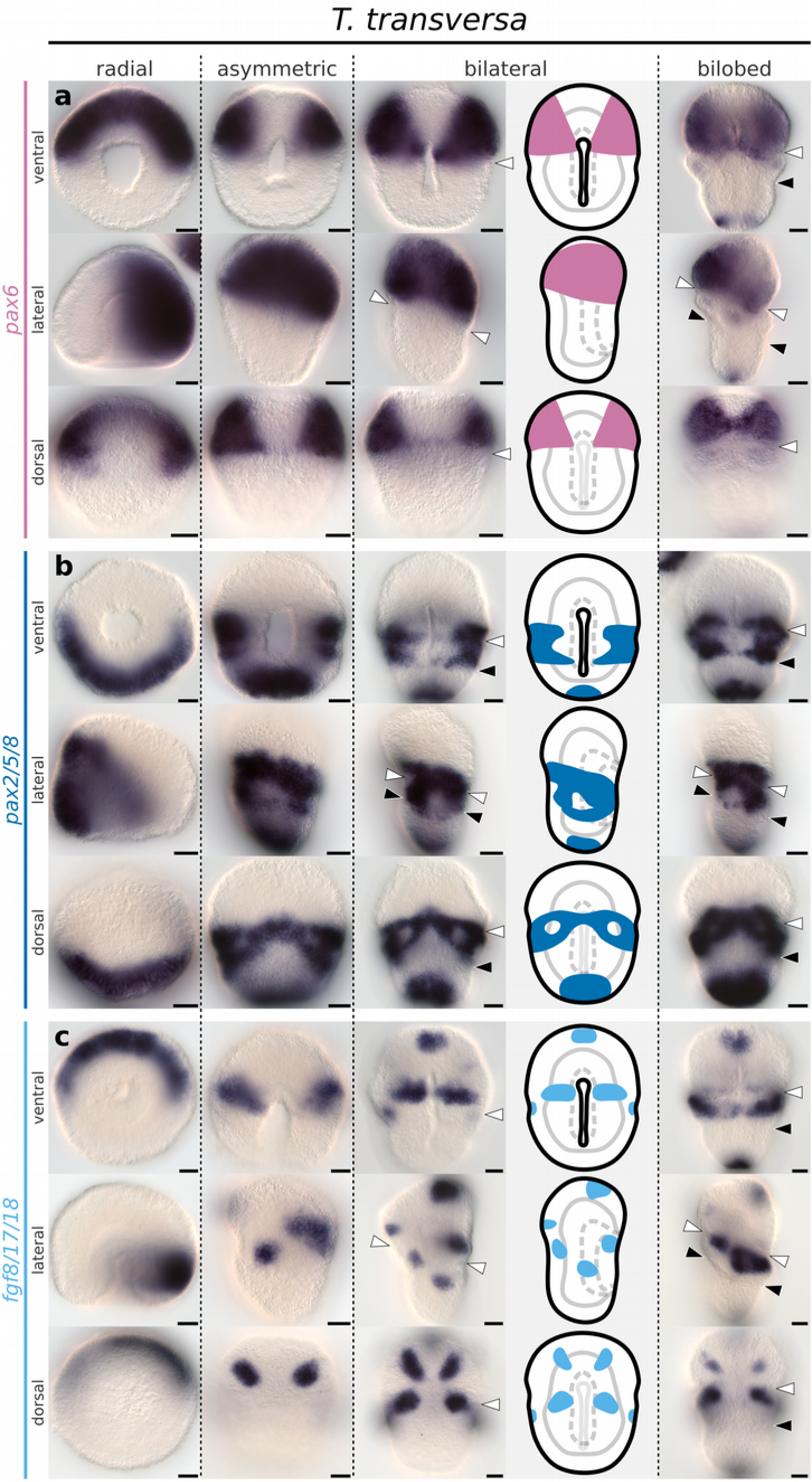
Whole mount in situ hybridization of *pax6*, *pax2/5/8* and *fgf8/17/18* orthologs in representative developmental stages of the brachiopod *T. transversa*. (a) Expression of *pax6*. (b) Expression of *pax2/5/8*. (c) Expression of *fgf8/17/18*. Anterior is top in all panels and ventral is to the right in all lateral views. Line drawings represent expression at the bilateral gastrula stage. White arrowheads mark the apical/mantle boundary and black arrowheads mark the mantle/pedicle boundary. Scale bars = 20 µm.

**Figure 7.**
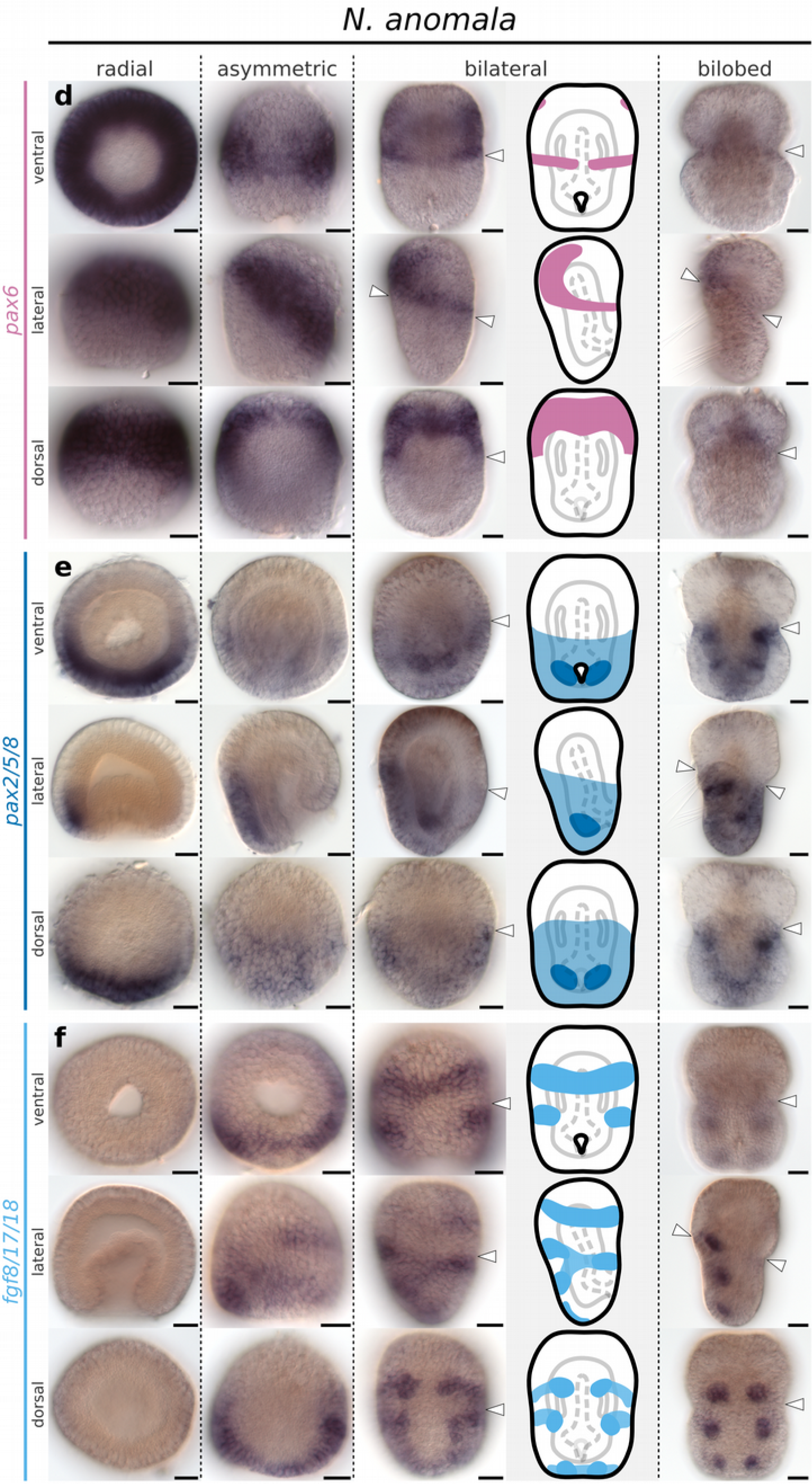
Whole mount in situ hybridization of *pax6*, *pax2/5/8* and *fgf8/17/18* orthologs in representative developmental stages of the brachiopod *N. anomala*. (a) Expression of *pax6*. (b) Expression of *pax2/5/8*. (c) Expression of *fgf8/17/18*. Anterior is top in all panels and ventral is to the right in all lateral views. Line drawings represent expression at the bilateral gastrula stage. White arrowheads mark the apical/mantle boundary. Scale bars = 20 µm.

*T. transversa pax2/5/8* expression begins as a posterior half-ring in the radial gastrula (Figure 6b) that overlaps with the *pax6* territory at the future apical/mantle boundary (Figure 8a). The transcripts cover the whole extension of the mantle ectoderm in the asymmetric gastrula, except for the chaetae sac primordia and blastoporal lips (Figure 6b), and contain the lateral domains of *en* (Figure 8d). The anterior limit of *pax2/5/8* expression crosses the apical/mantle furrow (Figure 6b) and overlaps with *pax6* expression at the posteriormost region of the apical lobe (Figure 8b–c). We corroborated this anterior limit of *pax2/5/8* transcripts by the combined analysis with *en* expression (Figure 8e–f); see Supplementary Fig. S7 for additional data. In *N. anomala*, *pax2/5/8* is expressed in a posterodorsal territory in the mantle lobe, limited by the apical lobe (Figure 7b). Different than *T. transversa*, we observe mesodermal expression of *pax2/5/8* in two pairs of domains between the chaetae sacs of *N. anomala* (Figure 7b). In summary, *pax6* and *pax2/5/8* are expressed in complementary domains, with a narrow overlap, that encircle the whole embryo before the lateral patches of *en* form transverse stripes. Thus, the apical/mantle furrow in both species is demarcated by the posterior limit of *pax6* expression with abutting domains of *en* in the mantle lobe.

**Figure 8.**
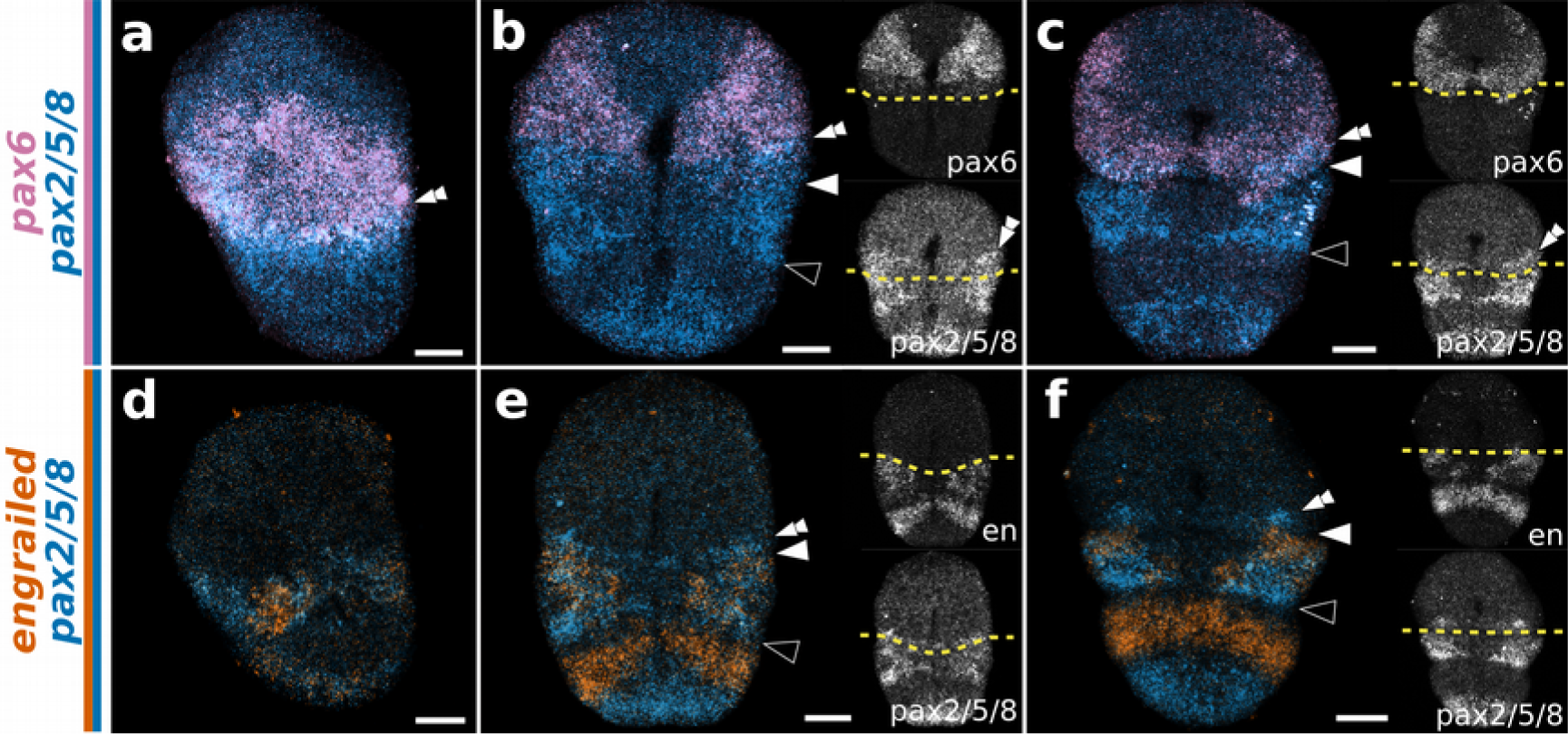
Whole mount double fluorescent in situ hybridization of *pax6* + *pax2/5/8* (a–c) and *en* + *pax2/5/8* (d–f) in the brachiopod *T. transversa*. (a,d) Asymmetric gastrula in lateral view. (b,e) Bilateral gastrula in ventral view. (c,f) Bilobed larva in ventral view. Anterior is top in all panels and ventral is to the right in all lateral views. Side panels show the relation between gene expression and the apical/mantle boundary (striped yellow line). White arrowheads mark the apical/mantle boundary and black arrowheads mark the mantle/pedicle boundary. Scale bars = 20 µm.

Transcripts of *fgf8/17/18* form a pair of transverse ventral bands in the apical lobe of *T. transversa* asymmetric gastrula (Figure 6c). These bands do not border the stripes of *en* posterior to the furrow (Supplementary Fig. S7). In addition, developing chaetae sacs express *fgf8/17/18* (Figure 6c) and are cleared from *en* and *pax2/5/8* expression (Supplementary Fig. S7). In *N. anomala*, *fgf8/17/18* expression initiates in the asymmetric gastrula with a similar anterior ventral band and a posterior domain encircling blastopore (Figure 7c). This posterior demarcates the anterior-most region of the mantle lobe at the bilateral gastrula but fades in the bilobed larva (Figure 7c). Similarly to *T. transversa*, chaetae primordia express *fgf8/17/18* (Figure 7c). Thus, the expression of *fgf8/17/18* is mainly associated with the chaetae sacs in both species and does not show a clear correlation with ectodermal boundaries.

### Over-activation of the Wnt pathway in *T. transversa*

The pervading role of Wnt signaling in the axial patterning of metazoans^46^, led us to investigate the role of the canonical Wnt pathway in the development of *T. transversa*. We found that the over-activation of Wnt signaling prevents the formation of the apical/mantle furrow (Supplementary Fig. S8). The posterior larval structures are expanded, the mantle lobe is not formed, and the expression of *en*, *wnt1* and *pax6* shifts more anteriorly (Supplementary Fig. S9). Early treatments entirely abolish the anterior domains of *en* and *wnt1*, but do not suppress the expression of *pax6* (Supplementary Fig. S9).

## Discussion

The evolutionary lineages of *T. transversa* (Rhynchonelliformea) and *N. anomala* (Craniiformea) diverged at least 500 million years ago^47^. It has been hypothesized that planktotrophy is ancestral for the Brachiopoda^48^ and that larvae of rhynchonelliforms and craniiforms might have evolved a lecithotrophic mode of development independently^49,50^. However, there are no extant planktotrophic larvae in the rhynchonelliforms, and for this reason, it is not possible to infer if the planktotrophy found in the larvae of extant linguliforms—the sister group of the craniiforms—is ancestral or derived. In our study, we compared species from the rhynchonelliforms and craniiforms to determine which traits are conserved or derived. We found that the trilobed larva of *T. transversa* and the bilobed larva of *N. anomala* share an apical/mantle boundary with a conserved ectodermal furrow morphology. A similar ectodermal furrow also delimits the apical lobe of the planktotrophic larva of *Lingula anatina*^51^, suggesting the apical/mantle boundary is an ancestral feature of brachiopod embryogenesis.

We found that *T. transversa* and *N. anomala* larvae show surprisingly consistent patterns of gene expression. Most expression domains in a species have a reciprocal, similarly positioned territory in the other species. In the trilobed larva of *T. transversa*, for example, *en* is expressed in a pair of anterior domains that correlate closely with the apical/mantle boundary, and in a pair of posterior domains bordering the mantle/pedicle boundary. The bilobed larva of *N. anomala* not only exhibits the correspondent *en* domains lining its apical/mantle boundary, but also the reciprocal posterior pair. We do not interpret these domains in *N. anomala* as a vestigial expression of *en*. First, in known cases of vestigial expression, an incipient aspect of the morphology is still present^52,53^. But there is no morphological evidence of a differentiated pedicle lobe in *N. anomala*. Second, the evolutionary origin of the pedicle lobe remains unresolved and it is unclear if it is an ancestral feature of brachiopod larvae. Nonetheless, the consistent expression patterns between *T. transversa* and *N. anomala* allow us to suggest that the presence of two pairs of lateral *en* domains is the ancestral condition for brachiopods, independent of the trilobed or bilobed larval morphology.

In order to better understand the evolution of *en* expression across bilaterians, we compared the brachiopod data with the available data from other animals (Figure 9). We found that most domains correlate to some kind of epithelial boundary such as segment boundaries of annelids, shell borders in molluscs, arthropod segment boundaries, seam cells in nematodes, collar/trunk boundary in hemichordates, brain boundaries in vertebrates and somite boundaries in cephalochordates (Supplementary Table S1). A closer look reveals that at its earliest instance of embryonic expression, most groups have an ectodermal expression of *en* in laterodorsal domains (Figure 9 and Supplementary Table S1). In annelids and brachiopods, a pair of lateral *en* domains are located at the head/trunk and apical/mantle boundary, respectively. In *D. melanogaster* the first stripe of *en* occurs at the cephalic furrow^54^ and in hemichordates *en* is expressed at the collar/trunk boundary^55^. This comparative data suggests that *en* expression is mainly ectodermal and might have been associated to the interface between an anterior (*e.g.* head) and a posterior region of the embryo (*e.g.* trunk), a division shared by most protostomes and deuterostomes.

**Figure 9.**
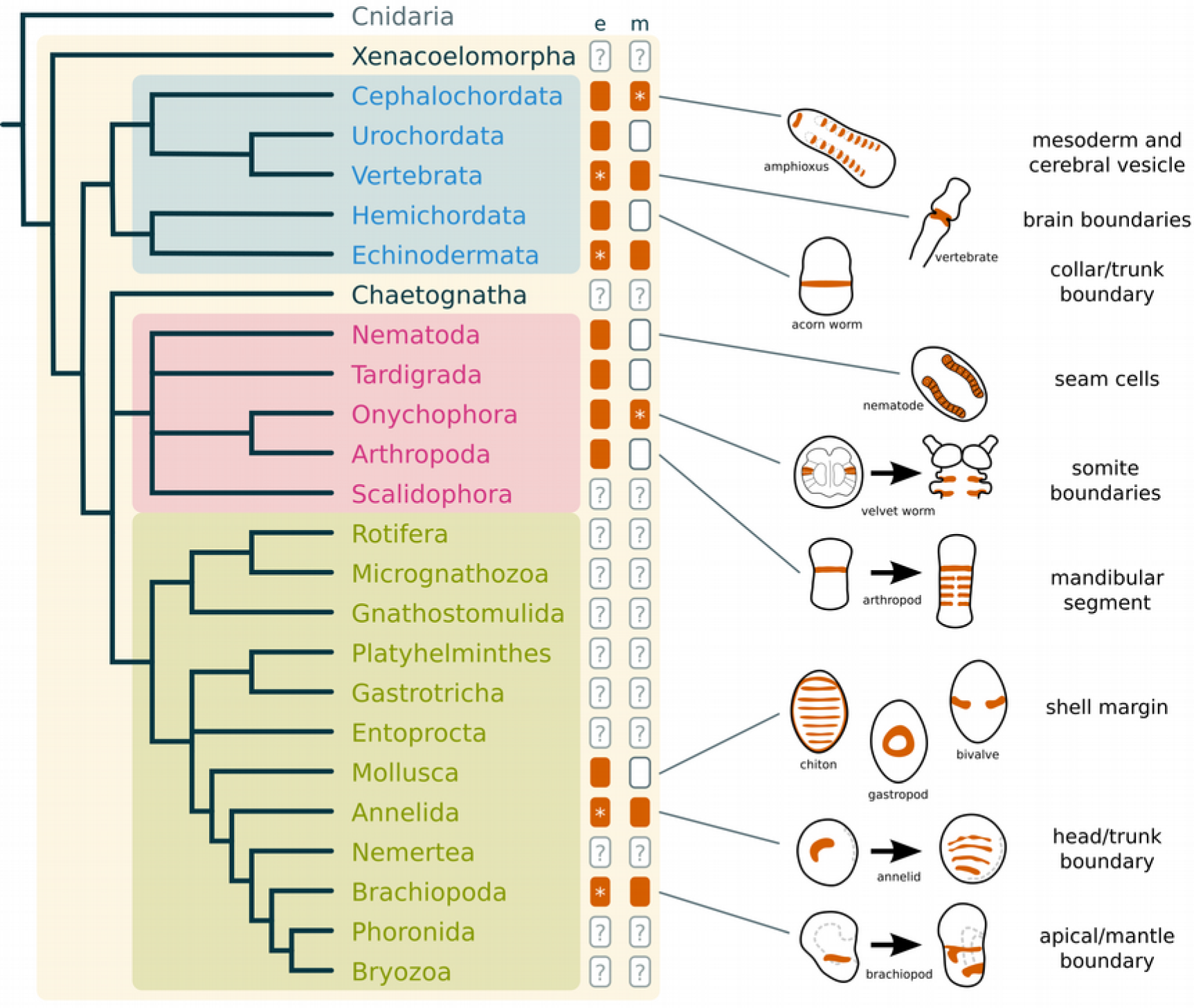
Comparative *en* expression in bilaterians. Phylogenetic tree based on recent data^29,60,61,74^ and gene expression compiled on the Supplementary Table S1. Boxes filled with color (vermillion) indicate the presence of *en* expression, either in the ectoderm (left column labeled with “e”) or mesoderm (right column labeled with “m”). If *en* is expressed in both ectoderm and mesoderm, asterisks indicate the tissue where *en* is first expressed during embryogenesis. Empty boxes indicate the absence of *en* expression and boxes with a question mark (”?”) indicate groups where the expression of *en* is unknown. Drawings show the expression of *en* (vermillion) in embryonic stages of representative groups. Early and late expression of *en* was depicted for Onychophora, Arthropoda, Annelida and Brachiopoda. Structures associated with *en* expression in each group are listed on the right.

In addition, our data indicates that the early patterning of the apical/mantle boundary of brachiopods is characterized by the overlapping expression of *pax6* and *pax2/5/8* (Figure 10a). Interestingly, the complementary patterns of *pax6* and *pax2/5/8* expression during early embryogenesis occur in the ectoderm of hemichordates^55^, cephalochordates^56,57^ and in the neuroectoderm of vertebrates^43,44^ (Figure 10b), suggesting that deuterostomes share a similar arrangement of gene products along the anteroposterior axis^58^. No nervous structures have been described at the apical/mantle boundary of larval brachiopods, so far^40,59^. Thus, the similarity to the brachiopod pattern could indicate that such genes were also involved in the axial patterning of the last common protostome ancestor. Or simply, that they have been independently coopted to these structures. Unfortunately, data on the expression of *pax6* and *pax2/5/8*, specially during early embryogenesis, are still scarce outside arthropods (Supplementary Table S1). Thus, it remains to be determined if the expression patterns of these genes in brachiopods are ancestral to the protostomes.

**Figure 10.**
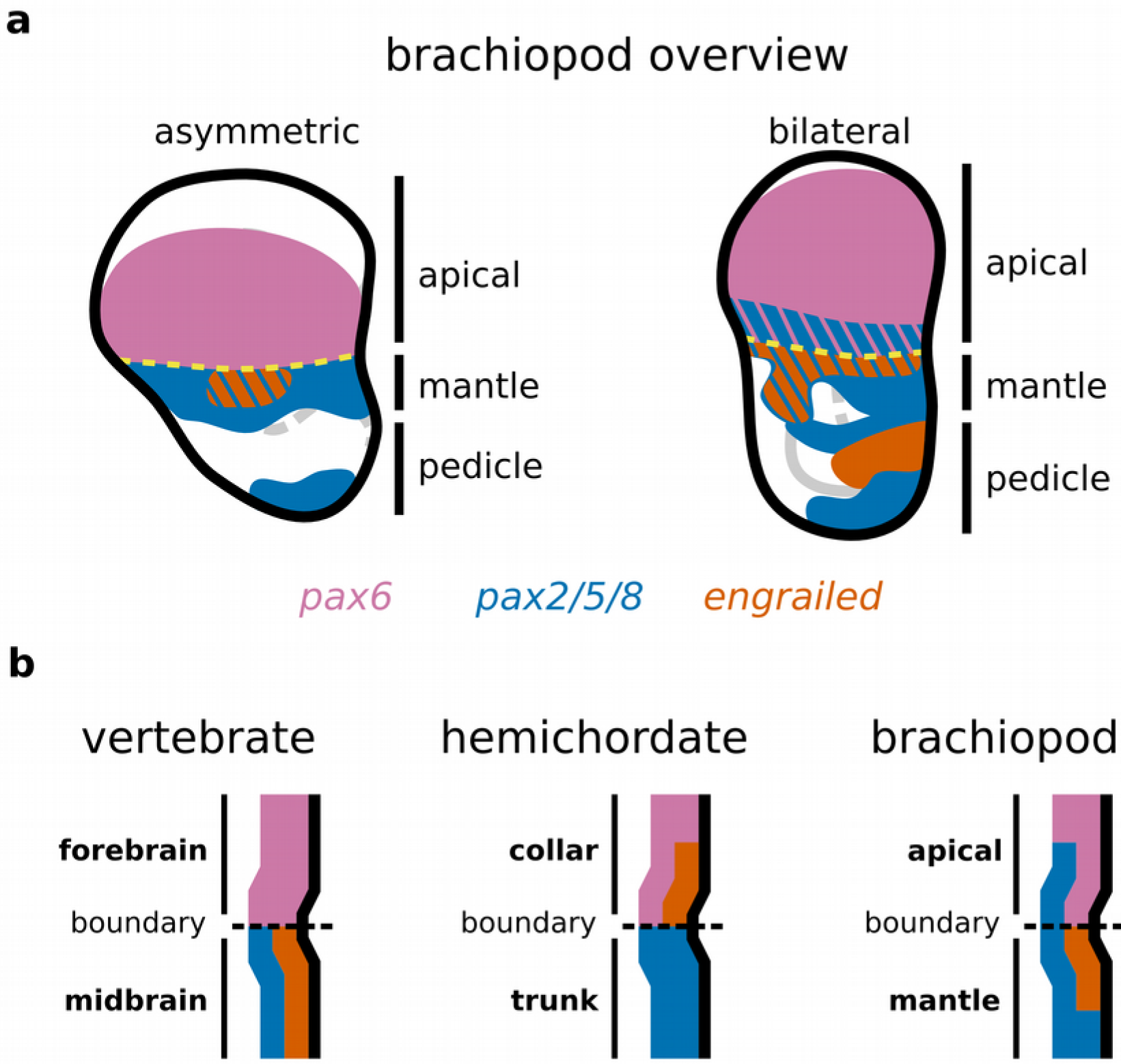
Summary of the brachiopod gene expression and the molecular profile of the apical/mantle boundary in comparison to other organisms. (a) Expression of genes related to the apical/mantle boundary with consistent expression between the brachiopods *T. transversa* and *N. anomala* (*pax6*, *pax2/5/8* and *en*). (b) Spatial relationship between the expression of *pax6*, *pax2/5/8* and *en* in the fore/midbrain boundary of the vertebrate brain (chicken)^43,44^, the collar/trunk boundary in the hemichordate (*Saccoglossus kowalevskii*)^55^ and the apical/mantle boundary in the brachiopod larval body (*T. transversa*).

Overall, the spatial arrangement and temporal expression patterns of our candidate genes suggest the apical/mantle boundary of brachiopods is not patterned by a segment polarity mechanism (Supplementary Fig. S10). Nevertheless, we show that abutting stripes of *en* and *wnt1* are not exclusive for the segment and parasegment boundaries of *P. dumerilii*^28^ and arthropods^7,19,24–26^, respectively—it also occurs in a non-segmental boundary of a larval brachiopod. Since the expression of segment polarity genes in annelids is variable and does not support a role in segment formation^5^, and recent phylogenies suggest the protostome ancestor did not have a segmented trunk-29,60, the segment polarity genes might be an arthropod-specific feature. Therefore, we interpret the adjacent expression of *en* and *wnt1* in *T. transversa*, *P. dumerilii* and arthropods as the independent recruitment of a common boundary patterning mechanism—not necessarily linked to segmentation—to these developmental boundaries. It remains yet to be found if annelids have a common set of genes patterning their homologous segments. Identifying such annelid-specific segmentation genes, and investigating their expression in other spiralians, will be essential to elucidate the evolution of segmentation mechanisms in the Spiralia, and how it compares to arthropod segmentation.

In conclusion, the conserved expression of *pax6*, *pax2/5/8* and *en* between *T. transversa* and *N. anomala*, and their correlation with the furrow morphology, suggests these genes might be involved in the patterning of the apical/mantle boundary of brachiopods. A broader comparison among bilaterians indicates the ancestral expression of *en* during early development was non-segmental and putatively related to the embryonic head/trunk boundary. We thus propose that *en* might have had a role in the early axial patterning of the protostome/deuterostome ancestor. Comparative expression data in the Xenacoelomorpha—the sister group to all remaining bilaterians^61^—as well as in less-studied protostome groups (*e.g.* gastrotrichs, rotifers, chaetognaths, nemerteans and priapulids) will be crucial to test this hypothesis.

## Materials and methods

### Sample collection

We collected adult brachiopods by dredging rocky ocean floor in Friday Harbor, USA (*T. transversa*) and Raunefjorden near Bergen, Norway (*N. anomala*) during reproductive season, December/January and September/October respectively. Fertilization of ripe gametes was conducted in the laboratory as described previously^37,62^ and the representative developmental stages were fixed in 4% formaldehyde for 1h, washed in PTw (1x PBS + 0.1% Tween-20) and kept in PTw at 4°C for antibody staining and in 100% methanol at −20°C for *in situ* hybridization.

### Immunohistochemistry

We permeabilized the embryos with several washes in PTx (1x PBS + 0.2% Triton X-100) for 2h and blocked with two washes of 1h in PTx + 0.1% BSA (Bovine Serum Albumin) succeeded by 1h incubation in PTx + 5% NGS (Normal Goat Serum). Samples were incubated with the primary antibodies (mouse anti-Tyrosinated Tubulin 1:500 and rabbit anti-Synapsin II 1:500) stored overnight at 4°C on a nutator. We removed the primary antibodies with three 5 min and four 30 min washes in PTx + 0.1% BSA, blocked in PTx + 5% NGS for 1h and incubated nutating overnight at 4°C with the secondary antibodies (Alexa Fluor 594 anti-mouse and Alexa Fluor 647 anti-rabbit 1:200). Secondary antibodies were removed with three 5 min followed by several washes in PTx + 0.1% BSA for 2h. We stained nuclei by incubating permeabilized embryos in DAPI 1:500 or Sytox Green 1:1000 for 2h. Nuclei staining was combined with f-actin staining by the addition of BODIPY FL Phallacidin 5 U/mL previously evaporated to remove methanol.

### Gene cloning and orthology

We identified orthologous genes by reciprocal BLAST searches using known sequences against the transcriptomes of *T. transversa* and *N. anomala*. We performed PCR gene specific primer pairs on cDNA of each brachiopod species synthesized with the SMARTer RACE cDNA Amplification kit (Clontech). We used RACE primers to clone *T. transversa en* and *pax6*. All other *T. transversa* and *N. anomala* genes were cloned with regular primer pairs (Supplementary Table S2).

Primers were designed with Primer3^63^. Orthology was assigned by aligning amino acid sequences of brachiopods against annotated genes using MAFFT 7.215^64^, retaining only informative portions of the alignment with GBlocks 0.91b with relaxed parameters^65^ and running a Maximum Likelihood phylogenetic analysis with RAxML 8.1.17^66^ using automatic model recognition and rapid bootstrap (Supplementary Fig. S11).

Alignments were manually verified using UGENE^67^. Resulting trees from the maximum likelihood analysis were rendered into cladograms using the ETE Toolkit^68^. Source files, alignments and scripts are available online at http://dx.doi.org/10.6084/m9.figshare.1473087.

### *In situ* hybridization

We synthesized antisense DIG-labeled riboprobes with MEGAscript kit (Ambion) and performed colorimetric *in situ* hybridization according to an established protocol^69^. Whole mount double fluorescent *in situ* hybridization was performed with the above protocol, but hybridizing the samples with a DIG-labeled and a DNP-labeled riboprobes. Samples were first incubated overnight at 4°C with Anti-DIG-POD conjugate diluted 1:250 in blocking buffer. After PTw washes, we developed the reaction with the TSA reagent kit Cy3 (Perkin Elmer). POD activity was inactivated by incubating 45 min in 0.1% H_2_O_2_ in PTw at room temperature followed by a 15 min incubation at 67°C in a detergent solution (50% formamide, 2x SSC, 1% SDS) and incubated overnight at 4°C with Anti-DNP-POD conjugate diluted 1:100 in blocking buffer. Second probe was developed in the same manner with the TSA reagent kit Cy5 (PerkinElmer).

### Imaging and image processing

Specimens were mounted in 80% Glycerol in PBS, 97% 2,2’-Thiodiethanol^70,71^ or Murray’s Clear (2:1 benzyl benzoate and benzyl alcohol solution) after a quick dehydration series in isopropanol (70%, 85%, 95%, 100%). After colorimetric *in situ* hybridization we imaged samples with a Zeiss AxioCam HRc mounted on a Zeiss Axioscope A1 using differential interference contrast technique (Nomarski). Fluorescent in situ hybridization and immunostainings were imaged in a Confocal Leica TCS SP5 and the resulting confocal stacks were processed in Fiji^72^. LUTs are available at http://github.com/nelas/color-blind-luts. We adjusted the levels of the final panels to improve the contrast using Fiji for confocal stacks and GIMP for photomicrographs. We created vector graphics and assembled the figure plates using Inkscape.

### Inhibitor experiments with 1-azakenpaullone

We sampled developing embryos of *T. transversa* from wild type cultures and incubated with a final concentration of 1 and 10 µM 1-azakenpaullone^73^ diluted in seawater. Embryos were picked at the mid-blastula and radial gastrula stage and fixed for immunohistochemistry and *in situ* hybridization at the bilateral gastrula and trilobed larval stage. Controls were treated with the highest concentration of dimethyl sulfoxide (DMSO) contained in the experimental samples (1% in seawater).

## Acknowledgements

We thank Chema Martín-Durán for crucial discussions and help with the in situ hybridization experiments, Carmen Andrikou and Kevin Pang for improving the manuscript, Yale Passamaneck, Yvonne Müller, Jonas Bengtsen, Daniel Thiel and Anlaug Boddington for the help with the collections, and Aina Børve for the laboratory guidance. We are thankful to the staff of Friday Harbor Labs and Espeland Marine Biological Station for their logistic support with the brachiopod collections. We also thank Alessandro Minelli for the comments on an earlier draft and two anonymous reviewers for the constructive feedback. The study was funded by the core budget of the Sars Centre and received support from the Meltzer Research Fund.

## Author contributions

BCV and AH designed the study and collected the material. BCV performed the experiments and analyzed the data. BCV and AH and wrote the manuscript. Both authors read and approved the final manuscript.

